# Generation of Monophasic Action Potentials and Intermediate Forms

**DOI:** 10.1101/2020.01.10.901793

**Authors:** Shahriar Iravanian, Ilija Uzelac, Conner Herndon, Jonathan J Langberg, Flavio H Fenton

**Affiliations:** Emory University, Atlanta, Ga; Georgia Tech, Atlanta, Ga

## Abstract

The Monophasic Action Potential (MAP) is a near replica of the transmembrane potential recorded when an electrode is pushed firmly against cardiac tissue. Despite its many practical uses, the mechanism of MAP signal generation and the reason it is so different from unipolar recordings is not completely known and is a matter of controversy. It is hypothesized that partial depolarization of the cells directly underneath the electrode contributes to the generation of MAP signals. In this paper, we describe a parametric, semi-quantitative method to generate realistic MAP and intermediate forms – multiphasic electrograms different from an ideal MAP – that does not require the partial depolarization hypothesis. The key ideas of our method are the formation of junctional spaces, i.e., electrically isolated pockets between the surface of an electrode and tissue, and the presence of a complex network of passive components that acts as a high-pass filter to distort the signal that reaches the recording amplifier. The passive network is formed by the interaction between the passive tissue properties and the double-layer capacitance of electrodes. We show that it is possible to generate different electrograms by the change of the model parameters and that both the MAP and intermediate forms reside on a continuum of signals. Our model helps to decipher the mechanisms of signal generation and can lead to a better design for electrodes, recording amplifiers, and experimental setups.

**SIGNIFICANCE:** Recording the Monophasic Action Potential (MAP) is potentially very useful in both experimental and clinical cardiac electrophysiology and can provide valuable information about the repolarization phase of the action potential. However, despite its benefits, it currently has only a small and niche role. The main challenge is the technical difficulties of recording an ideal MAP. Our results provide a better understanding of the mechanisms of the generation of cardiac electrograms and may help to optimize experiments and improve tools to achieve the full potentials of recording the MAP signals.

## INTRODUCTION

An extracellular electrode pushed firmly against cardiac tissue may record a near-perfect replica of the Transmembrane Action Potential (TAP) (1). This extracellularly-recorded intracellular potential is called the Monophasic Action Potential (MAP). The MAP is different from the low-amplitude and multiphasic signals recorded when an electrode is placed near cardiac tissue but does not press against it. Assuming the recording electrode is the positive input to the amplifier and a distant electrode the negative input, a MAP signal is positive, has a higher amplitude, and closely tracks the contour and duration of the TAP.

The characteristics of MAP have been known since the 1880s (2). Initially, the recordings were performed after causing injury to myocardium; however, atraumatic recording methods were found in 1930s (3), and further developed into clinical MAP catheters by Franz et al in the 1980s and 90s (4).

Despite its long history, the mechanism of MAP signal generation and the reason it is so different from other forms of unipolar recordings is not completely known and is a matter of controversy (5). It is hypothesized that pressing the electrode into the tissue partially depolarizes the cells directly underneath the electrode, probably through opening non-specific pressure-sensitive channels, and this contributes to the generation of MAP signals (2, 6). In this paper, we will show that such partial depolarization is not needed and that a natural interaction between the electrode and tissue can generate the MAP.

### Intermediate Forms

Figure 1 shows a group of signals collected by silver (Ag) electrodes (1.6 mm in diameter), coated with a layer of silver-chloride (AgCl), from five arterially-perfused rabbit ventricles (see (7) for the experimental details). The heart motion was suppressed with the injection of the myosin-inhibitor *blebbistatin* (8). The ground was a large Ag/AgCl electrode located in the bath remote from the hearts. All experiments conform to the current Guide for Care and Use of Laboratory Animals (9) and approved by the Office of Research and Integrity Assurance at Georgia Tech.

**Figure 1:**
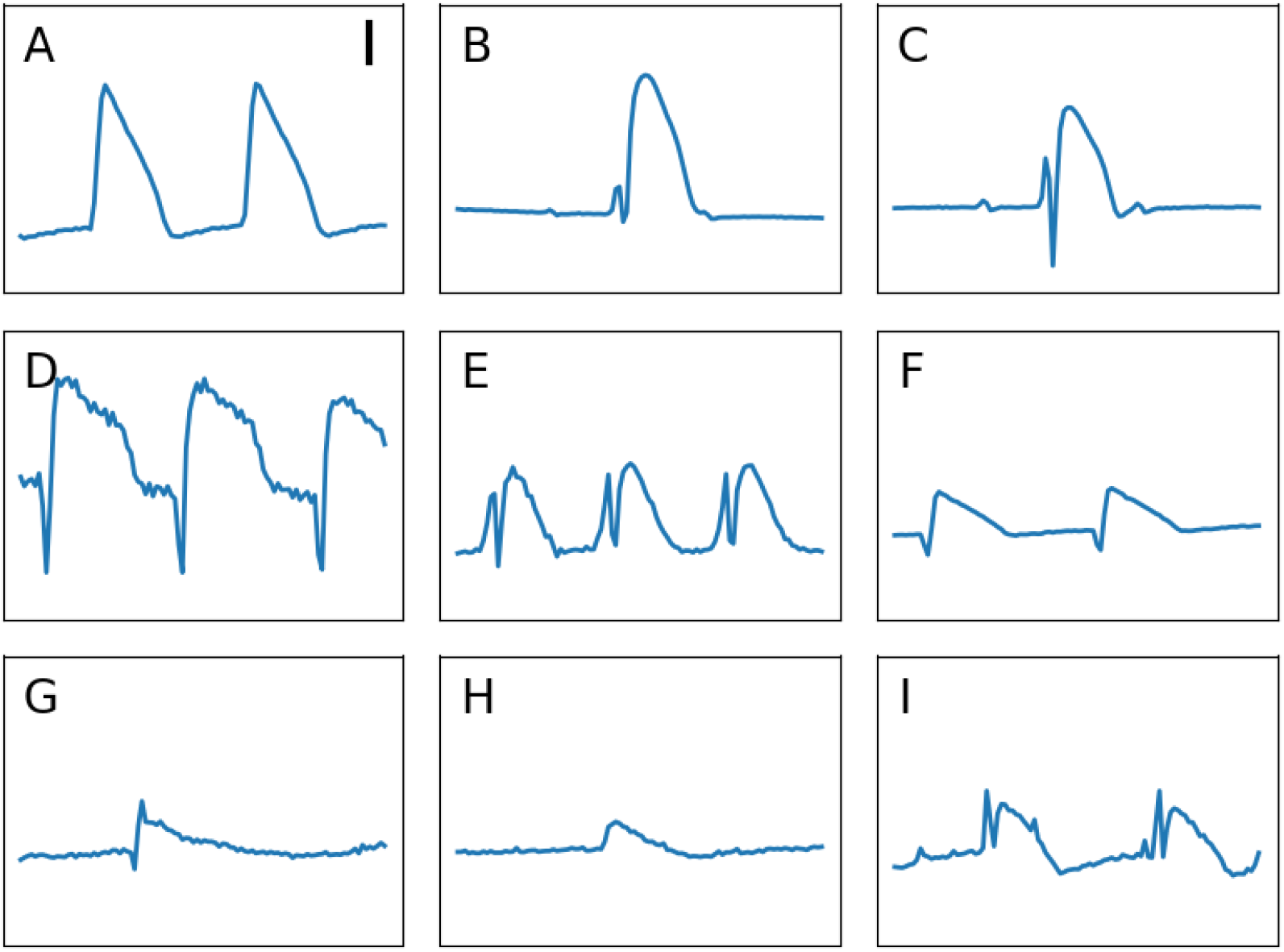
A collection of signals recorded using a Ag/AgCl electrode from five arterially-perfused rabbit ventricles on a Langendorff apparatus (**A**/**F**, **B**/**C**, **D**/**E**, and **G**/**H** pairs are from the same hearts). (**A**) and (**B**) are representative MAP; (**C-I**) are different intermediate forms. Note the sharp and multiphasic signals in the beginning of some the signal complexes. Each recording is 1 sec in duration. The vertical bar is 10 mV.

Only panels **A** and **B** depict acceptable MAPs; other panels exhibit various degrees of deformation and alteration. Usually, these *intermediate forms* are considered inadequate, and although useful for the timing of the upstroke of the action potential, their repolarization phase is discarded from further scrutiny. However, we believe that these forms contain useful information for deciphering the mechanisms of MAP generation. A key observation is that while pressing an electrode harder into the heart tissue, one sees a continuum of signals that gradually morph into MAP with no apparent threshold nor qualitative distinction between MAP and the rest of the intermediate forms. A realistic model should be able to generate both a typical MAP and various intermediate forms. In addition, the intermediate forms are the most frequently observed patterns in practice, and it would be helpful to be able to extract valid repolarization phase information from them.

## METHODS AND RESULTS

In this section, we describe a step-by-step process to simulate and model how electrodes record and shape cardiac electrical activity. Action potentials are the result of active membrane properties, namely the flow of currents through ion channels. However, cardiac tissue also possesses passive properties, including membrane capacitance and intracellular and extracellular resistance (10). Additionally, recording electrodes have passive properties (see below). The combination of all these passive elements can be modeled as a complex network of capacitors and resistors that shape and alter electrograms before they reach the recording amplifier. In this paper, our main goal is to develop a simple model of this passive network, which nevertheless can reproduce realistic signals, whether MAP or intermediate forms. For the sake of discussion, we split the model into two main components: biological (Bidomain Model) and non-biological (Recording Electrode); while acknowledging that the two are, in fact, intertwined and form a single network.

### Bidomain Model

The starting point in our modeling is the generation of extracellular potentials. We use a standard bidomain methodology (11, 12). For simplicity, we model a uniform one-dimensional piece of cardiac tissue (cable model); however, the main findings apply as well to 2D and 3D models. The governing equations are

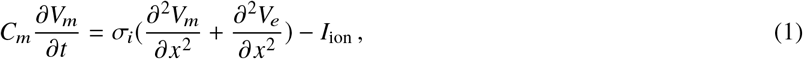

and,

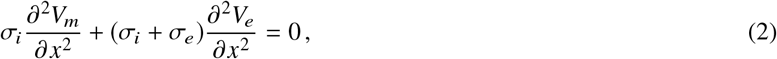

where *x* is the one-dimensional spatial coordinate, *V*_*e*_ is the extracellular potential, *V*_*m*_ is the transmembrane potential, *C*_*m*_ is the membrane capacitance (equals to 1 *μ*F/cm^2^), and σ_*e*_ and σ_*i*_ are, respectively, the extra- and intracellular conductance corrected for geometry. *I*_ion_ is the sum of membrane ionic currents. The intracellular potential, *V*_*i*_ = *V*_*e*_ + *V*_*m*_, is factored out from the formulation.

For the examples in this paper, we used the Mahajan rabbit ventricular ionic model (13). Again, the main findings are not sensitive to the details of the ionic models and any ventricular model, even a generic one such as the Beeler-Reuter model (14), should work. We chose this particular model because the recordings in Figure 1 were from rabbit ventricles.

Figure 2 shows a schematic of the bidomain model. The tissue is stimulated at 2 Hz, and the resulting *V*_*m*_ and *V*_*e*_, recorded from points **A** and **B**, are shown in Figure 3. *V*_*e*_ is a scaled down and inverted version of *V_m_*. This is an unsurprising and standard finding and is consistent with theoretical and analytic models (10, 15).

**Figure 2:**
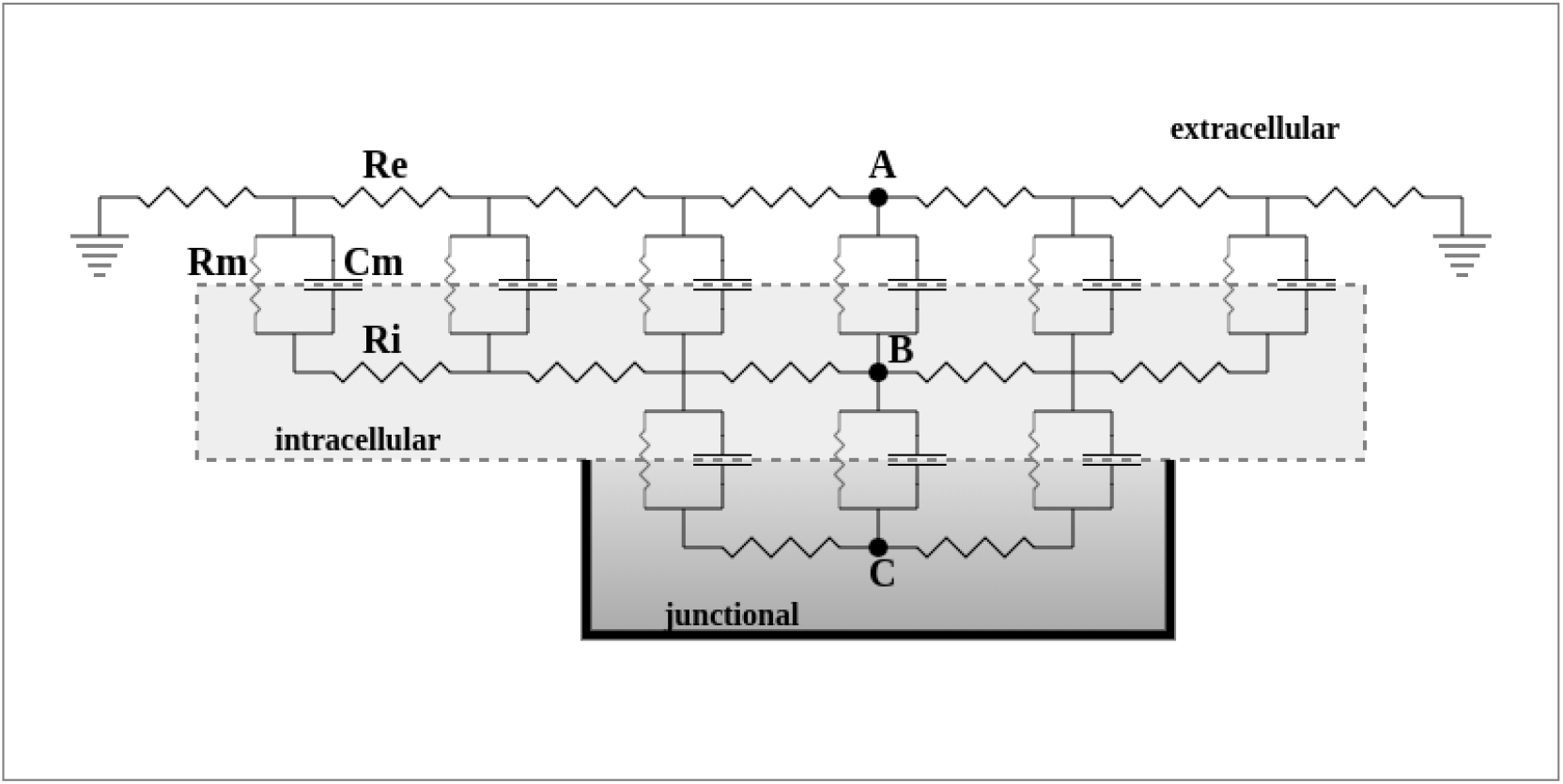
A schematic diagram of the double bidomain model. The intracellular, extracellular, and junctional domains are discretized as resistor networks. The connection between domains is through resistor and capacitor units of the cardiac membrane (**Rm** and **Cm**). Note that the extracellular domain is grounded, but the junctional space (the region in contact with the recording electrode) is not.

**Figure 3:**
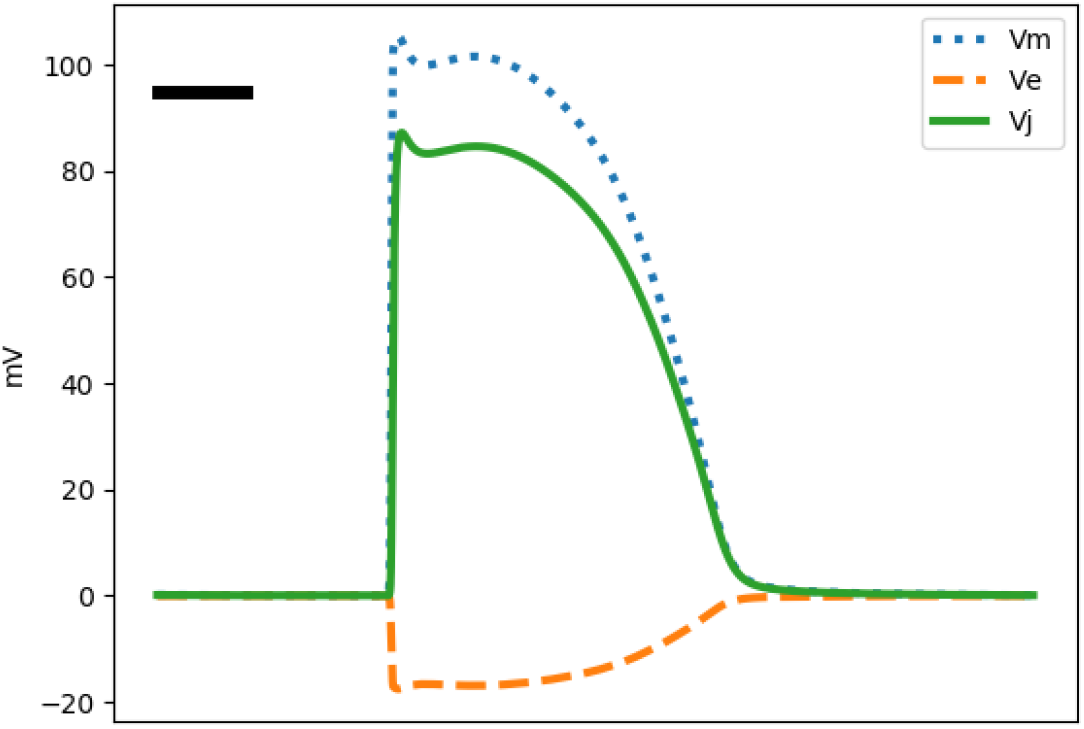
A representative transmembrane (*V*_*m*_), junctional (*V*_*j*_) and extracellular signals (*V*_*e*_) calculated using the double bidomain model for the Mahajan rabbit ventricular ionic model and assuming σ_*e*_/σ_*i*_ = 5. The signals are shifted vertically such that the potential is zero in diastole. Note that *V*_*j*_ ≈ *V*_*m*_ + *V*_*e*_. In addition, we have *V*_*e*_/*V*_*j*_ ≈ −σ_*i*_/σ_*e*_. The horizontal bar represents 50 ms.

#### Junctional Space

We hypothesized that an electrode pushing against the cardiac tissue sequesters some portion of the extracellular space. These *junctional* spaces (or domains) are located between the surface of the electrode and tissue and are isolated from the rest of the extracellular space (Figure 2). We assume that these spaces are small and, to the first approximation, do not affect the behavior of the cardiac cells, in contrast to the putative partial depolarization under electrodes (2, 6).

We model the potential inside a junctional space using the bidomain methodology. The main difference from modeling *V*_*e*_ is the boundary condition. For the junctional space, the Neumann (no-flux) boundary condition is imposed to enforce the fact that there is no direct current path between the junctional spaces and the bath. For a standard bidomain model, the boundary condition for the extracellular space is set as the Dirichlet condition to allow for the presence of ground connections in the bath or blood pool.

Changing the boundary condition has a dramatic effect on the potential. Figure 3 shows a representative junctional potential (*V*_*j*_) recorded from point **C**. The junctional space is essentially an extension of the intracellular space; therefore, *V*_*j*_ ≈ *V*_*i*_ = *V*_*m*_ + *V*_*e*_.

#### Signal Composition

If the entirety of an electrode surface is in contact with the junctional spaces, it will record a perfect MAP. This situation may apply to a suction catheter (16). However, most electrodes record a mixture of the junctional and extracellular potentials. Let *μ*, for 0 ≤ *μ* ≤ 1, be the *mixing ratio*. Then, the resulting composite potential, *W*, is *μV*_*j*_ + (1 − *μ*)*V*_*e*_. Here, *μ* is one of the parameters of the model.

Moreover, electrodes have a spatial extent. The junctional and extracellular spaces near an electrode are physically distinct. This spatial separation, in combination with the finite velocity of the depolarization wavefront, may introduce a time delay *τ* between the junctional and extracellular signals. *τ* is the second model parameter (a third one, *η*, will be introduced in the next section). Considering an electrode extent of 1-10 mm and conduction velocity of ≈ 50 cm/s, *τ* is in order of a few milliseconds. It can be positive or negative, depending on the direction of the wavefront and the alignment of the electrode.

Combining the effects of *μ* and *τ*, we have,

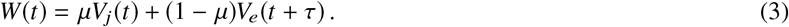

Figure 4 shows the resulting potential for a range of *μ* and *τ* values. For a small value of *μ* (*μ* = 0.1, the leftmost column), the dominant component of the signal is *V*_*e*_. As *μ* increases, the share of *V*_*j*_ also increases, and the output assumes more MAP features. The effect of *τ* is more pronounced on the upstroke phase that the repolarization phase, and the direction of the extra spike seen at the beginning of the signal complex.

**Figure 4:**
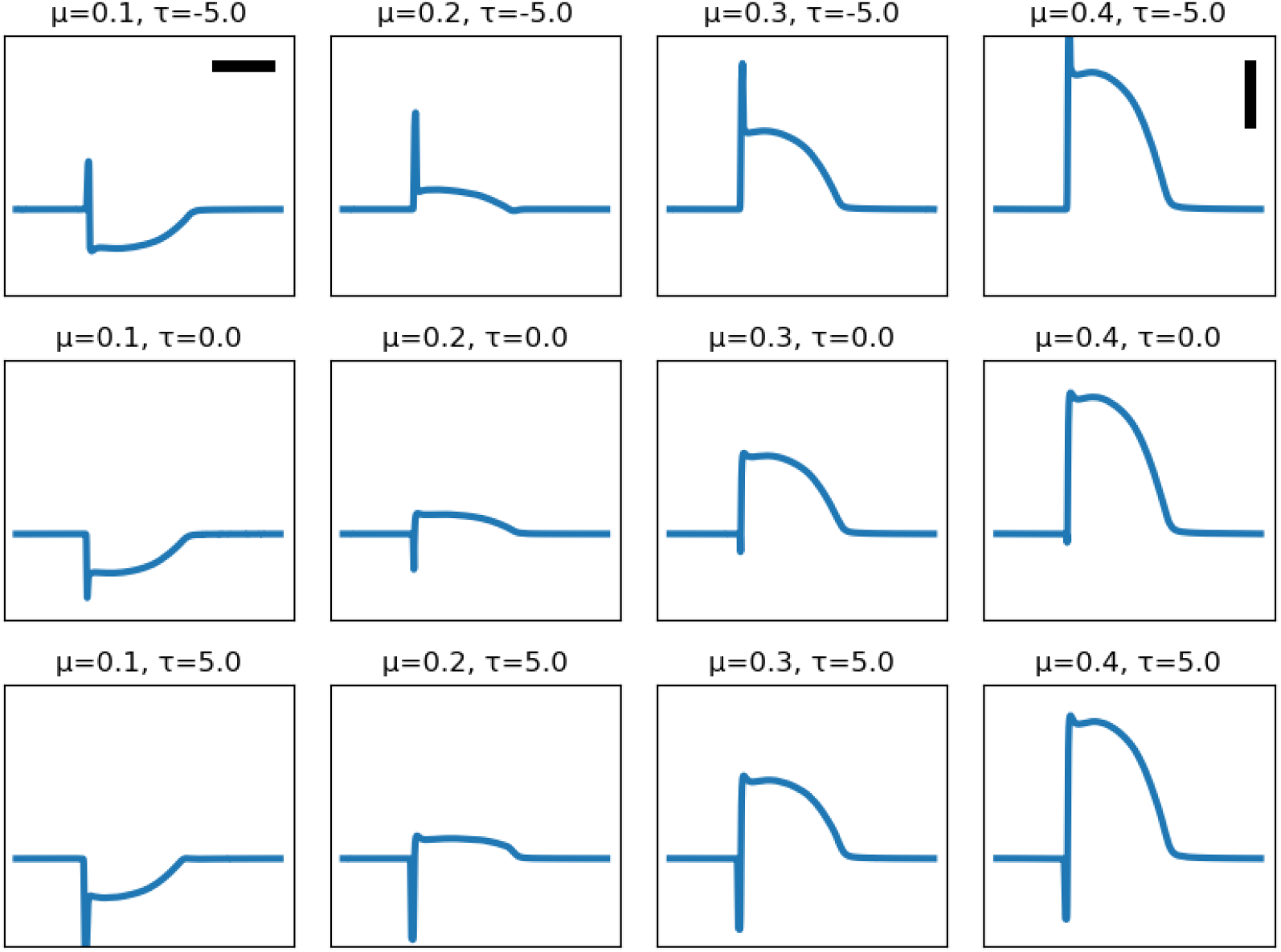
A collection of composite signals (**W**) for varying values of *μ* and *τ*. Note that as *μ* increases, the signals assumes more of MAP characteristics. The horizontal bar represents 100 ms and the vertical bar 10 mV.

The signals in Figure 4 represent what is presented to the recording electrode. If the electrode had ideal recording characteristics, then these signals also depict the input to the amplifier. However, an actual electrode tends to distort and modify the signal. This is the subject of the next section.

### Recording Electrode

#### Double-Layer Capacitance

Typical electrophysiology recording electrodes are either polarizing, made of an inert metal like platinum or iridium, or non-polarizing, usually made of AgCl coated Ag (7). We start the discussion with polarizing electrodes.

Charge carriers in conductors are electrons and in solution are ions. Transferring electrons to and from the solution requires a chemical reaction on the surface of the electrode, which inert metallic electrodes are incapable of doing (hence the name inert). Instead, charge transfer is achieved by charging and discharging a capacitor formed at the interface between the electrode and solution. This is called the double-layer capacitance (sometimes it is called the Helmholtz capacitor). This capacitance is in series with the resistance *R*_*s*_ of the current path in the tissue and bath to the return electrode (the ground).

We quantified the resulting RC circuit by using the Electrochemical Impedance Spectroscopy (EIS) methodology (17, 18). The technical details of the measurement system are described in (7). In summary, while the mechanical force was kept constant, the electrode was subjected to a set of pure tone sinusoidal currents. The resulting complex voltage response was measured, and the complex impedance, *Z*(*ω*), was calculated as the ratio of the measured voltage to the driving current. Here, *ω* = 2*π f* is the angular frequency. We used 20 test frequencies spaced evenly on a logarithmic scale between 2-2000 Hz, each applied for 5 seconds.

Figure 5A shows the impedance data for a typical clinical electrophysiology catheter (CRD-2, platinum electrode, area ≈ 6 mm^2^) (see the companion website to this paper, http://svtsim.com/eis, for a collection of impedance spectrograms for commonly used cardiac electrophysiology catheters and pacemaker leads).

**Figure 5:**
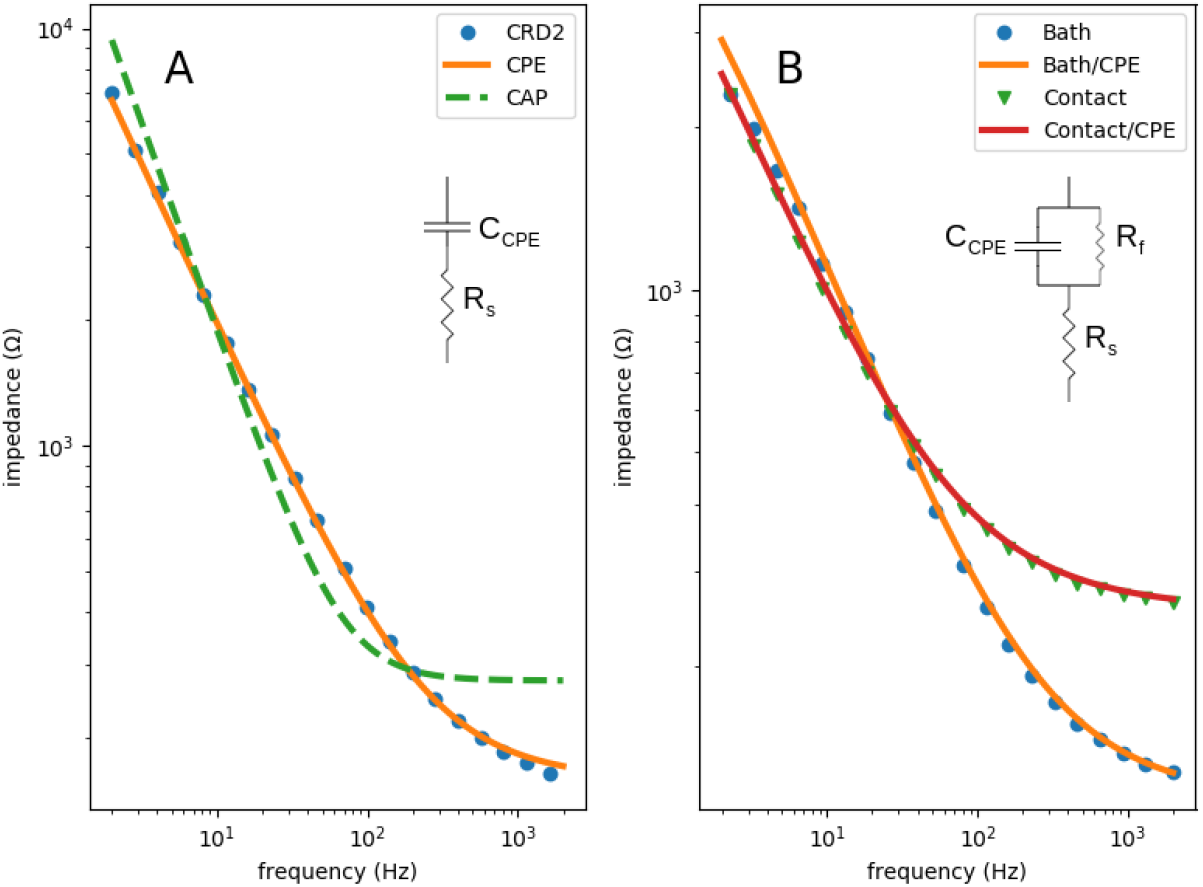
Impedance spectrograms of an inert metallic (**A**, CRD-2 electrophysiology catheter) and a partially AgCl coated Ag (**B**) electrodes. The spectrogram in **A** fits well with a CPE model (orange) but not an ideal capacitor model (dashed green). The best-fit is *Q* = 21 *μ*F/*s*^1−*α*^, *R*_*s*_ = 158 Ω, and *α* = 0.778. The spectrogram in **B** requires a parallel faradaic current resistor to fit well. The blue dots and the orange curve are for an electrode free in the bath (*Q* = 48 *μ*F/*s*^1−*α*^, *R*_*s*_ = 113 Ω, *α* = 0.709, and *R*_*f*_ = 8800 Ω.). The green triangles and the red curve are for an electrode pushing into the cardiac tissue (*Q* = 71 *μ*F/*s*^1−*α*^, *R*_*s*_ = 254 Ω, *α* = 0.675, and *R*_*f*_ = 16258 Ω.).

Our first attempt to fit the data to *Z*_*C*_ (*ω*) + *R*_*s*_, where

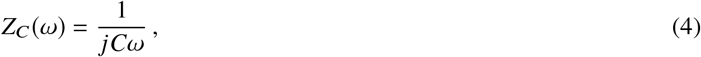

is the impedance of an ideal capacitor, *C* is capacitance, and 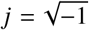 was not successful (Figure 5A, the green line). The reason is that the double-layer capacitor is not an ideal capacitor and exhibits frequency dependence. The double-layer capacitance is better approximated as a Constant Phase Element (CPE) (17–19). For a CPE, the impedance is defined as

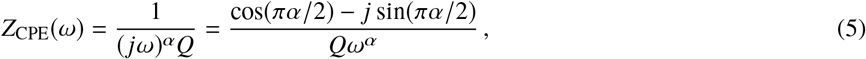

where *α*, for 0 ≤ *α* ≤ 1, is a parameter that measures the deviation of the CPE from an ideal capacitor (note that *α* = 1 corresponds to an ideal capacitor) and *Q* replaces *C* as a measure of the size of the capacitor. The orange curve in Figure 5A shows the result of fitting the experimental data to

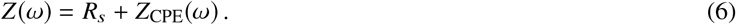

For the frequency range of interest in electrophysiology, the fit is excellent.

A non-polarizing electrode, such as a Ag/AgCl electrode, has a *faradaic* resistor (*R*_*f*_) in parallel to the double-layer capacitor (20). This current path is a result of the

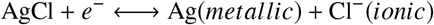

reaction on the surface of the electrode. Therefore, the consumption and regrowth of AgCl on the surface of the electrode allow for the exchange of electrons between the electrode and the ionic solution.

Figure 5B shows the impedance data for a Ag electrode partially covered with AgCl. Now, the total impedance is

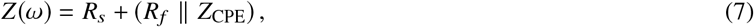

where ∥ stands for

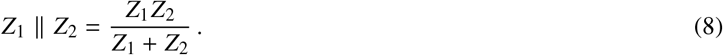

There are two sets of data in Figure 5B. In one, the electrode floats free in the bath (blue circles); in the other one, the electrode is pushing into the tissue (green triangles). Equation (7) fits well to both data sets.

#### Electrode as a High-Pass Filter

Armed with a reasonable model of the electrode surface, we can answer the question of signal distortion caused by an electrode. As mentioned in the previous section, a composite signal *W* is fed into the electrode. However, this is not what is sensed by the recording amplifier. It may seem that, considering the very high input impedance of modern amplifiers, the electrode impedance should be irrelevant. However, this is not the case! The main point is that the electrode is not a geometrical point. It has a nonzero surface area, such that different parts of the electrode surface are exposed to different potentials.

Figure 6 depicts a lumped-model of an electrode (**E**). This model is similar, but not identical, to models presented in (21) to describe signal distortion by metallic electrodes and glass microelectrodes, and in (22) to characterize EEG electrodes. The electrode is coupled to the tissue through a coupling capacitor (*C*_*w*_). The rest of the electrode is coupled to the bath/blood pool through the double layer capacitance (*C*_*s*_). In general, *C*_*w*_ ≪ *C*_*s*_; therefore, the impedance of *C*_*s*_ is essentially the same as *Z*(*ω*), where *Z*(*ω*) is calculated from Equations (6) or (7), depending on whether the electrode is polarizing or non-polarizing. We parameterize the coupling capacitor using *η*, for 0 ≤ *η* ≤ 1, such that *C*_*w*_ = *ηC*_*s*_, or equivalently,

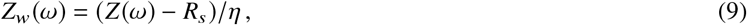

where *Z*_*w*_(*ω*) is the impedance of *C*_*w*_. As mentioned above, *η* joins *μ* and *τ* as the third parameter of our model.

**Figure 6:**
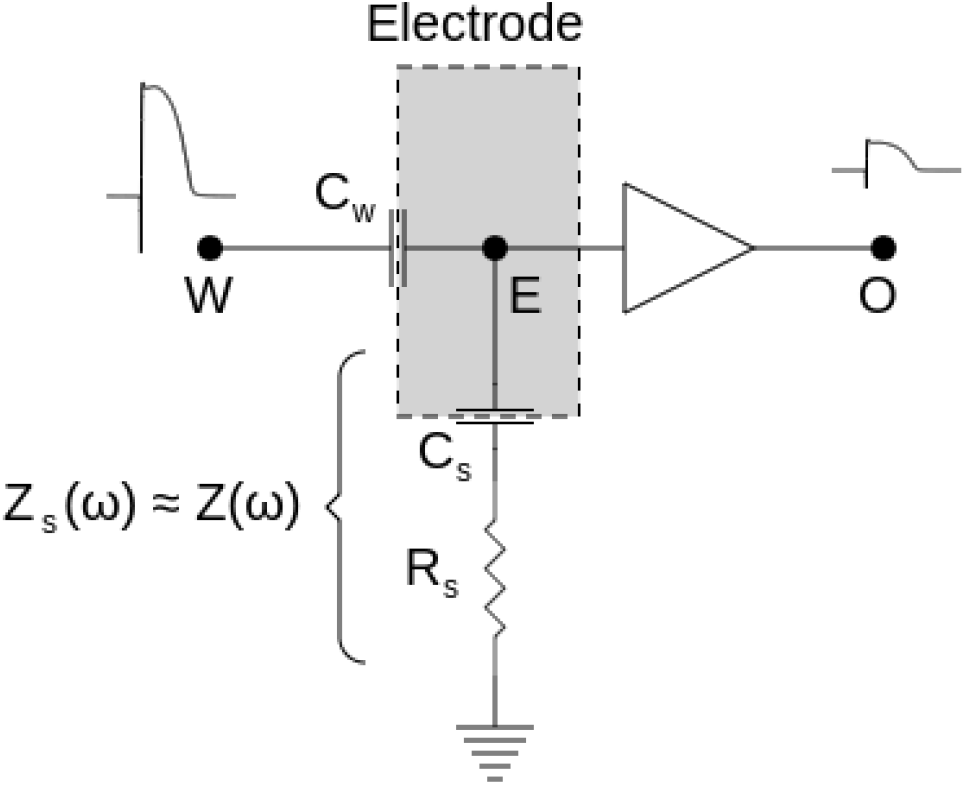
The lumped-element model of the electrode/tissue interface. The electrode surface is marked as a dashed rectangle. The surface impedance is split between a coupling capacitor (*C*_*w*_) to the source of the composite potential (*W*) and a capacitor connected to the ground (*C*_*s*_, assumed to be a CPE) through a resistor *R*_*s*_. The output of the amplifier is an attenuated and high-passed filtered version of *W*. The schematic is drawn for a polarizing electrode. For a non-polarizing electrodes, a faradaic resistor needs to be added parallel to *C*_*s*_.

Point **W** is the entry point of the composite signal *W*. The circuit in Figure 6 is a voltage-divider. Therefore, the transfer function becomes

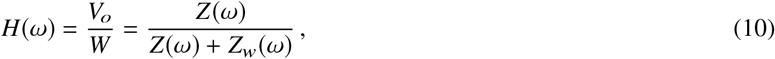

where *V*_*o*_ is the output potential. Substituting Equation 9, we have

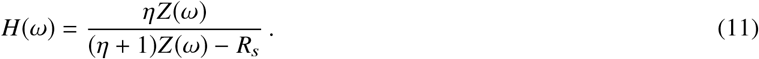

It should be noted that *Z*(*ω*) is complex; therefore, *H*(*ω*) is also a complex function and encodes both amplitude and phase information. At high frequencies, |*Z*| → *R*_*s*_; hence, |*H*| → 1. On the other hand, as *ω* → 0, |*Z*| becomes larger and |H| → *η*/(1 + *η*) ≈ *η* (remember that *η* is generally a small number). This is the behavior of a high-pass filter. However, the filter is leaky, such that even at the stopband (e.g., at DC) a portion *η* of the input signal still passes through the filter.

We can plug the values measured in Figure 5 into Equation 11 and calculate the frequency response of the corresponding electrodes (Figure 7). In the frequency range where the bulk of the signal power is (5-100 Hz), all these curves have a relatively flat response; hence, their primary effect is to attenuate the signal. This is one of the reasons why the amplitude of recorded signals is less than the amplitude of an action potential (~100 mV). The transfer function approaches 1 at higher frequencies; therefore, the electrodes pass the high-frequency portion of the signal (the upstroke of the action potential) with less attenuation. Algorithm 1 summarizes the process of estimating the recorded signal by an electrode with known characteristics. Applying it to the signals in Figure 4, we observe how the electrode preference for higher frequencies (high-pass filtering) alters the shape of the signal (Figure 8).

**Figure 7:**
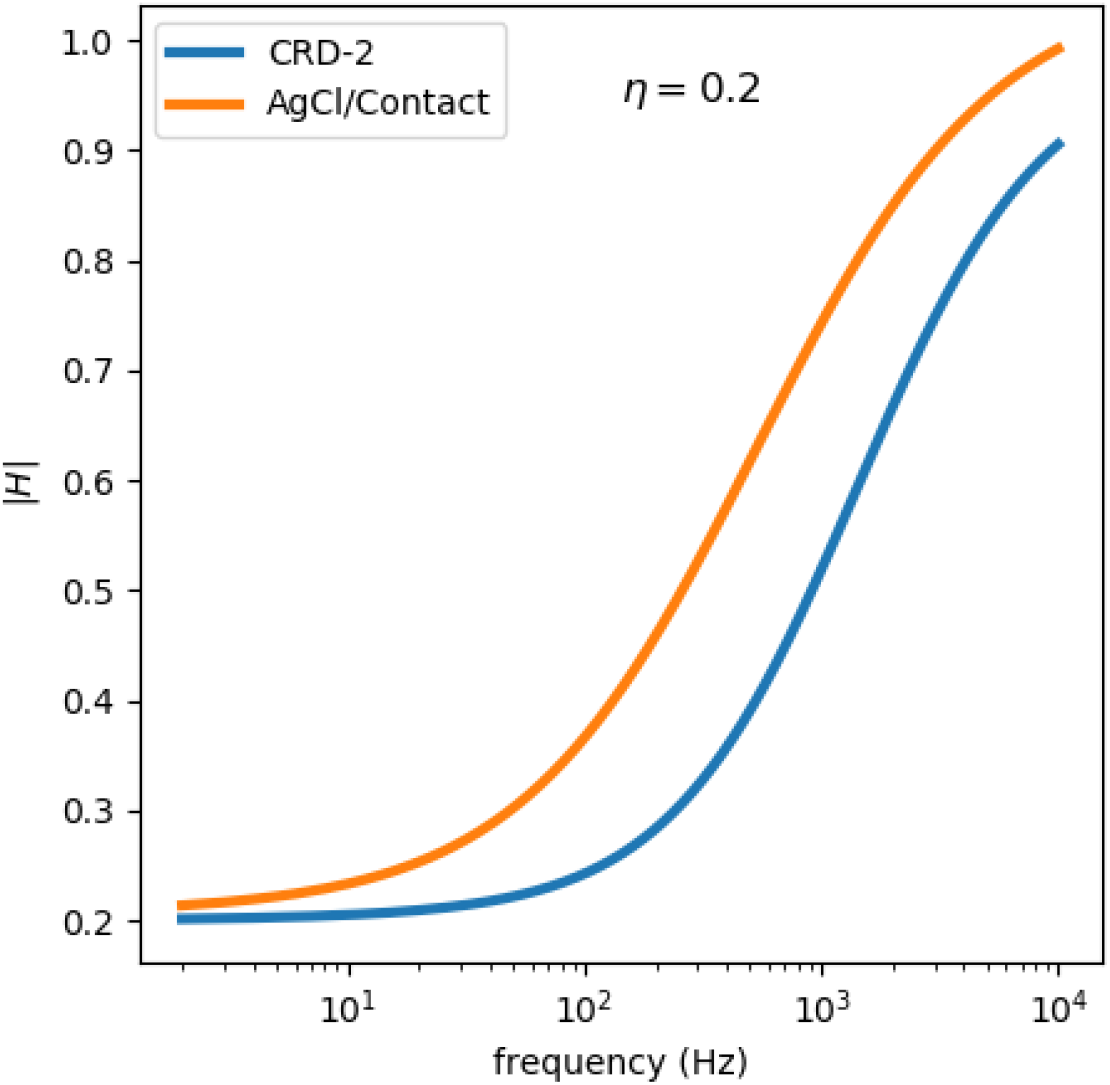
The transfer functions (the Bode magnitude graph) for two electrodes, a polarizing one (CRD-2) and a partially non-polarizing one (a Ag/AgCl electrode), assuming *η* = 0.2. Both curves depict high-pass filters with a non-zero response in the stopband. |*H*| stands for the magnitude of *H*(*ω*).

**Algorithm 1:**
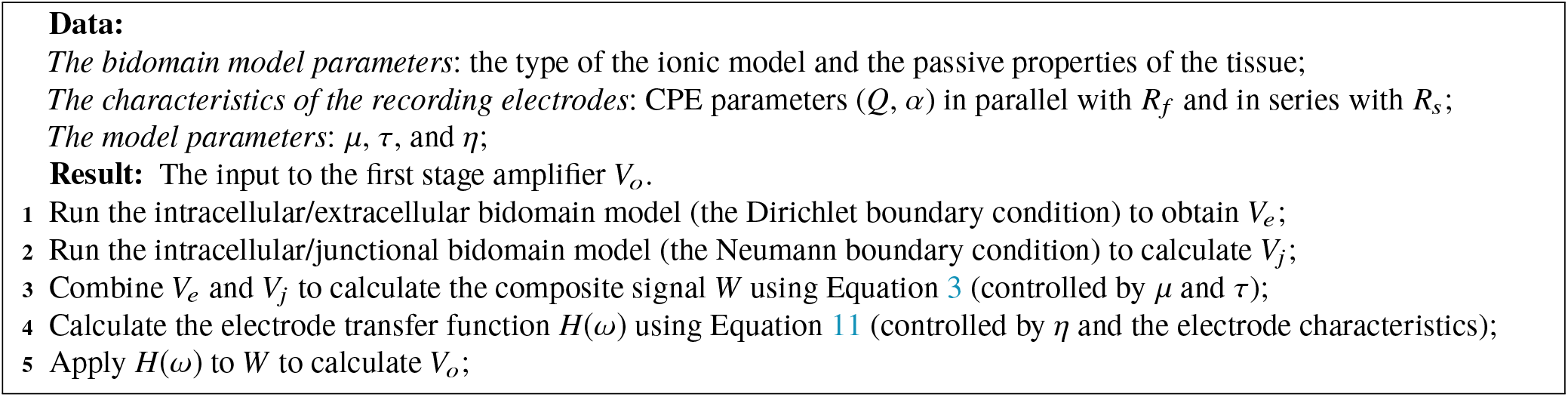
The summary of the algorithm to calculate the MAP and intermediate forms.

**Figure 8:**
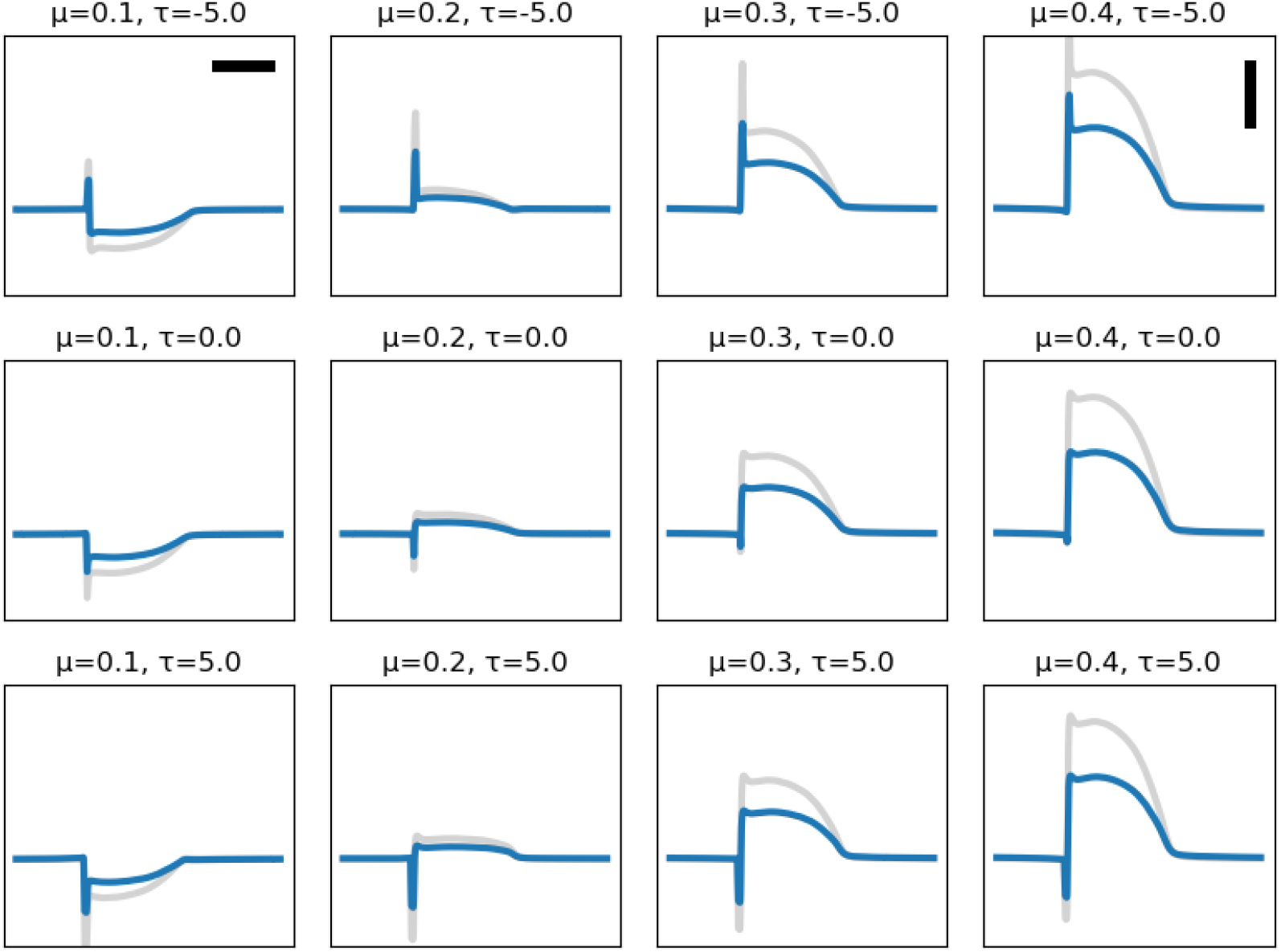
The effect of electrodes on shaping the signal. The figure shows the same signals as in Figure 4 (gray) subjected to the filtering effect of a polarizing electrode (the blue curve in Figure 7). Since the time-domain realization of a CPE is complicated, *H* was applied to *W* in the frequency domain. The horizontal bar represents 100 ms and the vertical bar 10 mV.

### Model Verification

The model presented above, especially Equations 10 and 11, is successful in reproducing realistic signals. However, this observation does not necessarily prove the correctness of the model. It is difficult to prove the model experimentally since *W* is hidden and hard to measure independently from the electrodes being modeled. One way to gain confidence in the soundness of the model is to observe that it produces the expected signal when the system is perturbed in certain ways. In this section, we present two such experiments to help with the verification of the model.

#### Dual Electrode

Figure 9A shows the schematic circuit used for the first experiment. We used a dual-electrode catheter. The tip electrode was very small (area 0.1 mm^2^) and in contact with the tissue (arterially-perfused guinea pig ventricles). The ring electrode was larger (area 5 mm^2^) and located in the bath. Both electrodes were made of a polarizing material (MP53N, a corrosion-resistant alloy commonly used in the medical field) and could be connected together electrically (via switch **S** is the schematic). The choice of material and the small size of the tip were by design, in order to exaggerate the expected signal distortion.

**Figure 9:**
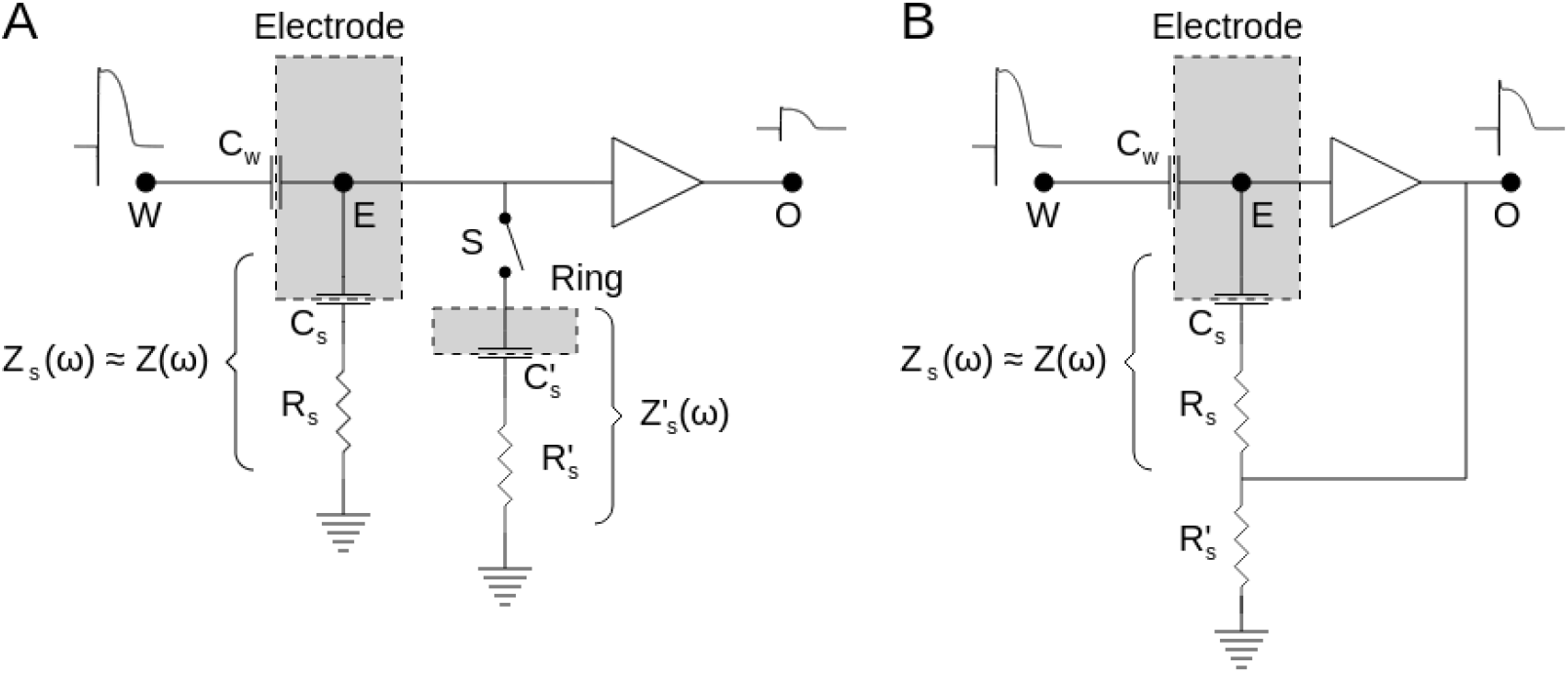
**A** The circuit diagram of the dual-electrode setup. The catheter has two electrodes. The tip is in contact with the tissue and is similar to Figure 6; whereas, the ring is in contact with the bath and can be turned on and off using switch **S**. **B** The bootstrapped version of the lumped-element model of the electrode/tissue interface. The output of the unity gain amplifier forms a positive feedback loop and boosts the ground connection of the voltage divider. We assume 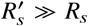.

Figure 10A shows the signals recorded from the tip (blue, **S** open) and from the combined tip and ring (orange, **S** closed). Note that the signal is attenuated and distorted after the switch was closed. As we will see below, we can analytically estimate the ratio of the transfer function of the tip-only configuration to the transfer function of the combined electrodes. The same ratio can be measured experimentally as the square root of the ratio of the estimated power spectra of the signals (Figures 10C and 10D, the purple dots).

**Figure 10:**
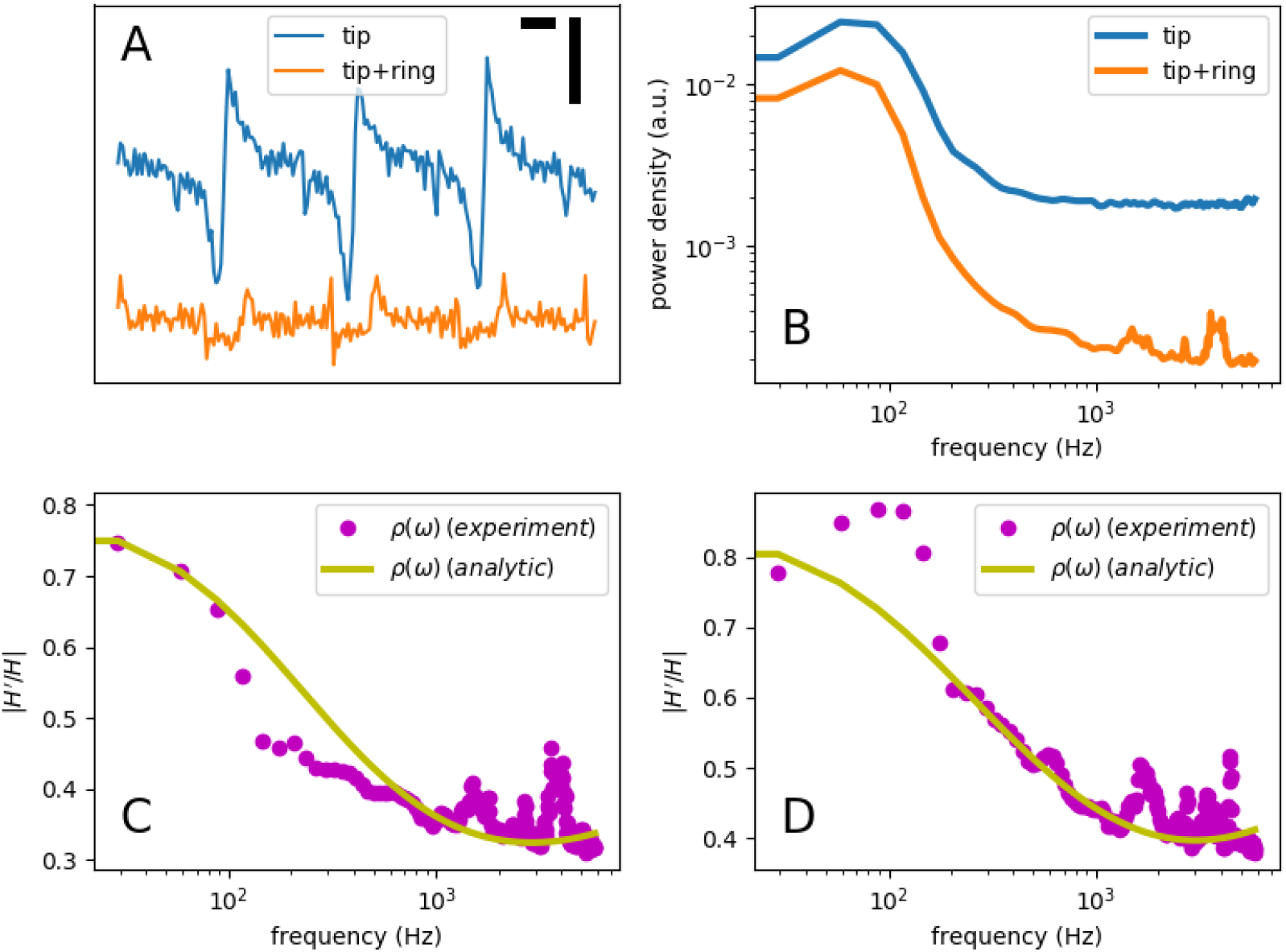
The experimental data recorded from two guinea pig ventricles using a dual-electrode catheter. **A** Recorded signals from the tip electrode with the ring off (blue) and on (orange). The horizontal bar represents 100 ms, and the vertical bar is 10 mV. Each recording was 20 sec long. **B** The power spectra of the signals shown in **A**. **C** The experimental (the purple dots, based on the data in **B**) and analytic (the yellow line, calculated using Equation 14) ratio of the transfer functions (the ring-on to ring-off ratio). **D** The experimental (purple) and analytic (yellow) ratio for another ventricle, similar to **C**.

Let *H*(*ω*) be the transfer function of the tip electrode (**S** open) and *H′*(*ω*) be the transfer function of the combined setup (**S** closed). Our goal is to find *ρ*(*ω*) = *H′*(*ω*)/*H*(*ω*). The starting point is Equation 10,

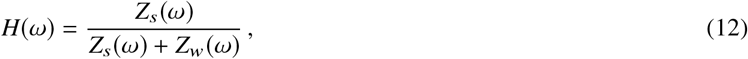

where *Z*(*ω*) is the complex impedance of the tip and *Z*_*s*_ (*ω*) is the impedance of the (unknown) coupling capacitor (from hereon, we will use *Z* instead of *Z*(*ω*) to reduce clutter). Similarly, for the combined electrode,

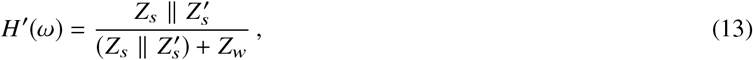

where *Z′*, is the impedance of the ring. Considering that 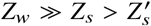, we have

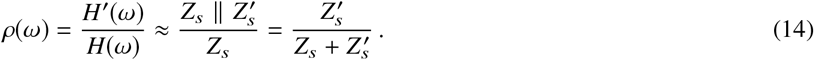

Using Equation 14, the analytic transfer function ratio (the yellow curve in Figure 10C and D) were calculated and were found to be a reasonable fit of the measured ratio (the purple dots).

#### Bootstrapping

Another modification of the system is shown in Figure 9B. Here, the output of the amplifier, with unity gain, is fed back as the ground electrode. In electronics, this is called *bootstrapping* and is used to convert a passive to an active filter.

With bootstrapping, the input to the electrode becomes *W* + *V*_*o*_ instead of *W*. Let *G*(*ω*) be the new transfer function (*H* (*ω*) is the original transfer function). The actual gain of the amplifier is *β*, which is assumed to be near 1. We have,

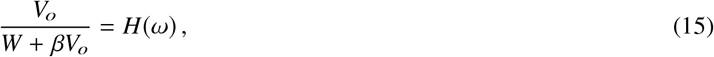

therefore,

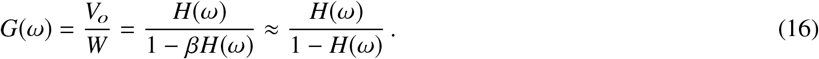

Figure 11A shows the transfer functions before and after bootstrapping for a AgCl electrode. The two curves mostly coincide in low frequency but diverge at high frequencies. The reason is that when *H* is small, 1 − *H* is near 1. As *H* becomes larger and approaches 1, the 1 − *H* term becomes smaller and boosts the resulting gain. Hence, *G*(*ω*) is an exaggerated high-pass filter compared to *H*(*ω*).

**Figure 11:**
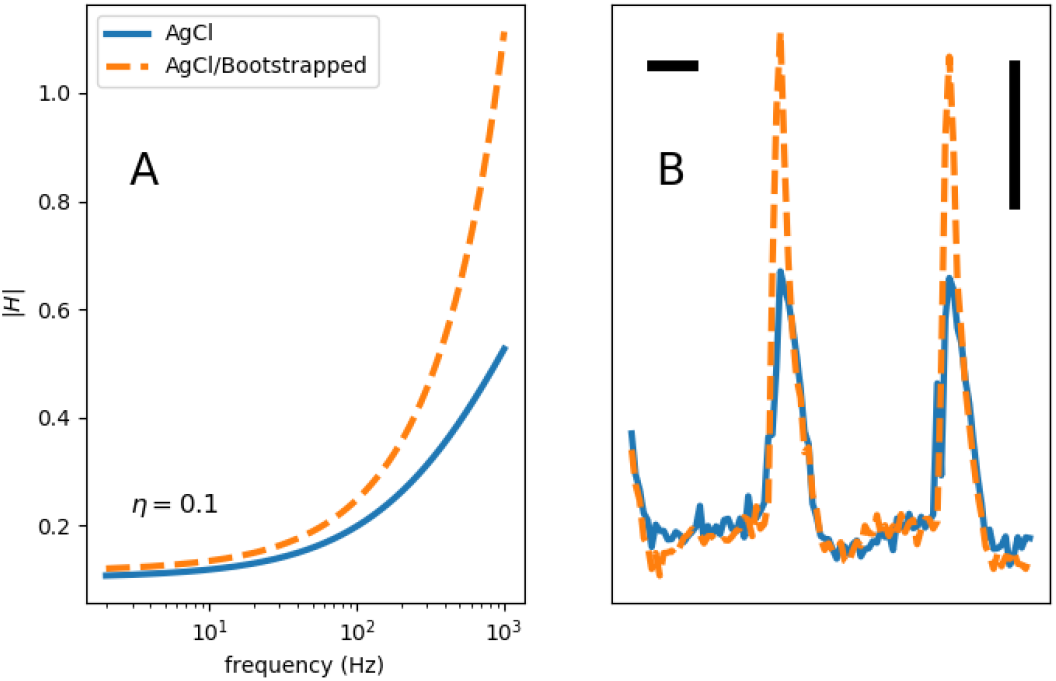
The effects of bootstrapping on the filter transfer function (**A**) and a representative experimentally-recorded signals from a guinea pig ventricle (**B**).

Figure 11B demonstrates the effect of bootstrapping in recordings from a guinea pig ventricle. The bootstrapped signal (orange) has a larger high-frequency component compared to the baseline recording (blue). This can be seen as the significant difference in the upstroke region (high frequency) compared to the essentially identical repolarization phase (low frequency).

## DISCUSSION

Recording cardiac electrical activity with the help of electrodes – including MAP catheters, microelectrodes, electrophysiology catheters, pacemaker leads, and other forms – is the cornerstone of both experimental and clinical electrophysiology. However, what is recorded may not be, and usually is not, a faithful reproduction of the electrical activity at the tissue level. The complex electrical network formed by the passive elements of the cardiac tissue and the passive properties of the electrode itself distorts the signal.

In this paper, we presented a methodical process to generate realistic signals. The core idea of our model is the presence of junctional spaces, i.e., electrically isolated pockets between the electrode surface and tissue. These spaces are similar to the isolated pockets of fluid seen in thin solid-liquid-solid films (23), and provide a window into the cardiac cells. Another important consideration is that electrodes are not geometrical points and have finite surface areas. This fact is relevant to both the mixing of junctional and extracellular potentials and the presence of a high-pass filter at the junction of an electrode and cardiac tissue created because of the double-layer capacitance. Importantly, there is no need to assume partial cellular depolarization as a result of the electrode force. We also showed that MAP and intermediate forms are qualitatively similar and can be generated by the same model with a continuous change of the parameters (i.e., *μ*, *τ*, *η* defined above).

Understanding the mechanisms of signal generation helps with a better design for electrodes, recording amplifier, and the experimental setup. For example, we can reduce the detrimental effects of an electrode on the amplitude and shape of the signals by making its response curve as flat as possible. Based on the transfer function equations (Equations 10 and 11), we can achieve this goal by either making the electrode impedance (*Z*_*w*_) smaller, making the series resistor (*R*_*s*_) larger, or increasing the electrode contact (*η*). The former can be done by either decreasing the faradaic shunt resistor (*R*_*f*_) by using an Ag/AgCl electrode or increasing the capacitance of the electrode, for example, by increasing its surface area as in fractal pacemaker leads (24–26). *R*_*f*_ can be made larger by pushing the electrode harder into the tissue, which also increases both *μ* and *η*, or by using a suction electrode. All these techniques are commonly used, and the reason they work can be understood within the unified framework presented in this paper.

Most electrophysiology practitioners, whether experimental or clinical, develop an intuitive feel for how the properties of electrodes affect the signal quality. For example, it is generally appreciated that a larger electrode produces a lower frequency signal. This observation is usually attributed to a larger “antenna” effect; i.e., a larger electrode surface area records from a larger volume of tissue and averages out the signal; hence, the lower frequency. However, intuition without a solid theoretical background can be misleading. As we saw, a more accurate explanation of the effects of a larger electrode is that all electrodes act as a high-pass filter, but a larger electrode has a larger capacitance and, therefore, the cutoff frequency of the filter is shifted toward lower frequencies.

Despite its benefits, recording MAP has only a small and niche role in clinical and experimental electrophysiology. The main challenge is the technical difficulties of recording an ideal MAP, which requires the experiment to be optimized toward recording MAP. The fact that MAP and intermediate forms are just different points on the same continuum means that we can settle for a less than ideal MAP and potentially correct for distortions based on the passive properties of the electrode.

## CONCLUSION

The MAP is one of the two extremes on the continuum of possible signals (electrograms) recorded by an electrode touching myocardium. The electrograms do not require force-dependent partial depolarization; instead, they originate from isolated junctional spaces and are shaped by the passive properties of the electrode and tissue.

## AUTHOR CONTRIBUTIONS

S.I. designed the research, and collected and analyzed the data. I.U., C.H., and F.F.H. performed the experiments. J.J.L. provided the concept. S.I. and J.L.L. wrote the manuscript with input from other authors.

## ACKNOWLEDGMENTS

The study was partly supported by NSF grant No. 1762553 and NIH grant No. 1R01HL143450-01.

